# Integrated Analysis of Cross-Links and Dead-End Peptides for Enhanced Interpretation of Quantitative XL-MS

**DOI:** 10.1101/2023.05.26.542474

**Authors:** Andrew Keller, Xiaoting Tang, James E. Bruce

## Abstract

XL-MS provides low-resolution structural information of proteins in cells and tissues. Combined with quantitation, it can identify changes in the interactome between samples, for example, control and drug-treated cells, or young and old mice. A difference can originate from protein conformational changes altering the solvent-accessible distance separating the cross-linked residues. Alternatively, a difference can result from conformational changes localized to the cross-linked residues, for example, altering the solvent exposure or reactivity of those residues or post-translational modifications on the cross-linked peptides. In this manner, cross-linking is sensitive to a variety of protein conformational features. Dead-end peptides are cross-links attached only at one end to a protein, the other terminus being hydrolyzed. As a result, changes in their abundance reflect only conformational changes localized to the attached residue. For this reason, analyzing both quantified cross-links and their corresponding dead-end peptides can help elucidate the likely conformational changes giving rise to observed differences of cross-link abundance. We describe analysis of dead-end peptides in the XLinkDB public cross-link database and, with quantified mitochondrial data isolated from failing heart versus healthy mice, show how a comparison of abundance ratios between cross-links and their corresponding dead-end peptides can be leveraged to reveal possible conformational explanations.

## Introduction

Chemical cross-linking with mass spectrometry (XL-MS) provides low resolution structural information about protein conformations and configurations by virtue of identified cross-linked residues being within the maximum span of the applied cross-linker^1–3^. Identifying a cross-linked peptide pair, however, requires that the attached residues on their proteins not only reside within the maximum cross-linker span but also be solvent accessible and reactive. Because this approach can be applied to cells *in vivo* in their natural environments, XL-MS can help elucidate diverse protein conformations in a variety of cells^4–7^, tissues^8^, and organs^9^.

Quantitative XL-MS can reveal how interactomes change in response to various conditions such as drug treatment, disease or aging, providing ratios of the abundance of identified cross-links in one sample relative to another^7,10–17^. Efforts in our lab first investigated quantitative cross-linking with applications to cellular phenotypic and pharmacological differences using SILAC, which revealed cross-linked peptide level changes that contributed to functional differences^7,13^. Technologies such as iqPIR^18,19^ greatly enhance the ability to quantify cross-links in an accurate and reproducible manner.^10^. When cross-link abundance differences are due to level changes of the cross-linked protein, one expects all cross-links derived from that protein to exhibit abundance level alterations similar to the protein. In contrast, observation of a cross-link abundance level change that differs from that of the protein(s) of origin may arise due to conformational or interaction differences of the cross-linked protein(s). These changes can alter the solvent-accessible distance between the cross-linked residues and thereby affect cross-link levels. However, such an observation could also be due to protein conformations with distinct solvent accessibility or reactivity of the cross-linked residues. For example, the existence of salt bridges at lysine residues in some but not all protein conformations can produce a conformational-dependence of cross-linker reactivity at these sites, even if the distance between sites is not conformationally-dependent^20^. Finally, because cross-linked peptide pairs are identified after tryptic cleavage of sample proteins, a change in PTMs at the cross-linked residues or elsewhere on the cross-linked peptides, including at enzyme cleavage sites adjacent to the cross-linker reactive sites, can affect the abundance of the cross-linked peptide pair.

Cross-linked samples often include dead-end (DE) peptides originating from the cross-linker attached to a protein residue at only one of its two reactive sites, the other being hydrolyzed. These are not true cross-links since they do not span two protein residues and cannot give information on protein configurations as maximum distances between protein residues like a cross-link. Nevertheless, DE peptides resultant from iqPIR reactions can be quantified in the same manner as iqPIR cross-linked peptides and in the past, were used merely to estimate proteome quantitation^21^. However, iqPIR DE peptide relative abundance levels in two samples is influenced by the same factors that affect the abundance of a cross-link discussed above, with the exception of conformational differences altering the solvent-accessible distance between two residues. DE peptide abundance levels reflect only conformational differences localized to the DE peptide itself and its linked residue. In fact, DE peptides provide information analogous to other covalent labeling techniques such as radical footprinting^22^ or HDX^23^, albeit with less depth, precision, or temporal dynamics than either of these methods. Here, we describe how a direct comparison between the relative abundance of cross-linked peptide pairs and their DE peptide counterparts in samples derived from animals with phenotypic differences can help elucidate the conformational causes of observed cross-link abundance differences, whether they involve an alteration in the solvent-accessible distance between the cross-linked residues or changes of other conformational features localized to one or both residues. We show how DE peptides can now be uploaded and displayed in the public cross-link database XLinkDB^24–27^ alongside their corresponding cross-links and how this integrative analysis helps identify likely causes of observed cross-link abundance level changes.

## Methods

### TAC versus Sham heart mitochondrial cross-link and DE data

Previously described heart mitochondrial quantitative XL-MS data obtained from 6 pairs of TAC versus Sham mice^28^ was used for this study. Raw data and search results are available at the ProteomeXchange Consortium via the PRIDE^29^ partner repository with the dataset identifier PXD027757. In addition, 16 early eluting strong cation exchange (SCX) fraction runs enriched for DE peptides that were not previously deposited are available with the dataset identifier PXD040864 (**Username**, **Password**: L3sCx0Ye).

### Quantitative data analysis

The combined DE peptide and cross-link data was quantified with iqPIR^21^. The data contained 6 pairs of TAC and Sham paired mice that generated 6 independent log2ratios of TAC versus Sham. All replicate pairs were combined together into a combined ratio that serves as an average. We filtered for log2ratios with a 95% confidence limit no greater than 0.5 and a minimum of 8 contributing quantified ions (peptide and/or fragment). We then required all cross-links to have quantitation in 4 or more of the replicates. Cross-links with log2ratios significantly different from those of both cross-linked proteins were obtained by a maximum Anova p-value of 0.05 applied only to the combined ratios.

### Correspondence between cross-links and DE peptides

Cross-links are associated with a DE peptide if and only if one of the cross-linked peptides has the same amino acid sequence, including modifications, as the DE peptide as well as the same site of attachment to the cross-linker based on its residual stump modification. This strict relationship ensures that the quantitation of the DE peptide truly encompasses all the localized environmental factors that will contribute to the abundance of the cross-link. Beyond the localized contributions represented by the corresponding DE peptides, the abundance of the cross-link also depends on the distance and solvent accessibility of the path between the cross-linked residues.

### Evaluating consistency of quantitation between cross-links, their corresponding DE peptides, and proteins to help infer possible explanations for cross-link abundance differences

iqPIR quantitation of cross-links and DE peptides yields a log2ratio of abundance in two compared samples such as TAC versus Sham mouse mitochondria, which is calculated by averaging the log2ratios of multiple quantified peptide and fragment ions in one or more MS2 spectra in which they are identified. Because many independent measurements contribute to these values, they are robust, accurate, and reproducible. Anova p-values are computed to assess whether a cross-link or DE peptide log2ratio is different from that of the protein(s) from which it originates. They are similarly used to assess whether a cross-link log2ratio is significantly different from that of either of its corresponding DE peptides. In this manner, cross-links with log2ratios significantly different from those of the cross-linked proteins (Anova p-value ≤ 0.05) were selected for closer inspection. Among them, the Anova p-values were used to find cases in which 0, 1, or both of the corresponding DE peptides had log2ratios similar to those of the cross-link (Anova p-value > 0.08) and significantly different from those of the originating protein (Anova p-value ≤ 0.05).

The quantitation of cross-links and corresponding DE peptides is used together to identify consistent possible conformational changes that cause cross-link abundance differences not due to protein levels, those altering the solvent-accessible distance between the cross-linked residues or those localized to a peptide, altering the accessibility or reactivity of its attached residue or PTMs on any of its residues. The directionality of proposed changes of conformational features (increase or decrease) depends on whether the cross-link log2ratio is greater or less than zero, as discussed below.

In cases in which none of the corresponding DE peptides have log2ratios significantly different from those of the originating protein, it is inferred that the abundance difference of the cross-link in the two compared samples is due to a conformational change altering the solvent-accessible distance between the cross-linked residues, including, in the case of an inter-protein cross-link, a change of interaction between the two proteins such as increasing or decreasing association if the log2ratio is greater or less than zero, respectively. When both ends of the cross-link are attached to the same protein and the cross-link is homodimeric, obtained from two distinct protein molecules, or if known structures of the protein contain a protein dimer, it is possible that the observed abundance difference is due to a change of dimerization in the compared samples. These possible explanations are displayed in the hypothesis column to the right of the log2ratio in the XLinkDB dataset table.

When a DE peptide corresponding to a cross-link has a log2ratio that is significantly different from its originating protein yet similar to that of the cross-link, the cause of the observed cross-link abundance difference is inferred to include a conformational change localized to the DE peptide such as increased or decreased solvent accessibility or reactivity of the attached residue, if the log2ratio is greater or less than zero, respectively. In addition, decreased or increased levels of PTMs reported by dbPTM^31,32^ to occur on residues of the peptide or its preceding amino acid are also included as possible explanations for log2ratios greater or less than zero, respectively, since they would reduce the levels of the identified peptide lacking those PTMs. If the cross-linked peptide itself contains PTMs, then increases or decreases in their levels are proposed for log2ratios greater or less than zero, respectively. These possible explanations are displayed in the hypothesis column.

XLinkDB supports any method of quantifying cross-links and DE peptides^30^. As long as the log2ratios are accompanied by their standard deviations and number of replicate measurements, the Anova p-values described above can be computed to enable the display of possible explanations for cross-link abundance differences not due to protein level change in the hypothesis column.

## Results and Discussion

DE peptides can now be uploaded and displayed in XLinkDB alongside cross-links, including looplink peptides containing both residues to which a cross-link is attached. When data are quantitative, whereby each cross-link is assigned a log2ratio reflecting its relative abundance in two samples, the quantitation of the cross-linked peptides and corresponding DE peptides— those with the same peptide sequence and cross-linked amino acid—can be displayed together in a heatmap (Figure 1). One can easily compare the abundance ratio of a cross-link with those of the corresponding DE peptides to help infer possible differences in protein conformations and configurations giving rise to them. Details of iqPIR quantitation, including the MS2 spectra of the contributing peptide and fragment ions, can be viewed by clicking on the cross-link log2ratio or on the DE peptide heatmap panels below. On XLinkDB, one can filter to view only cross-links with at least one or both corresponding DE peptides.

**Figure 1.**
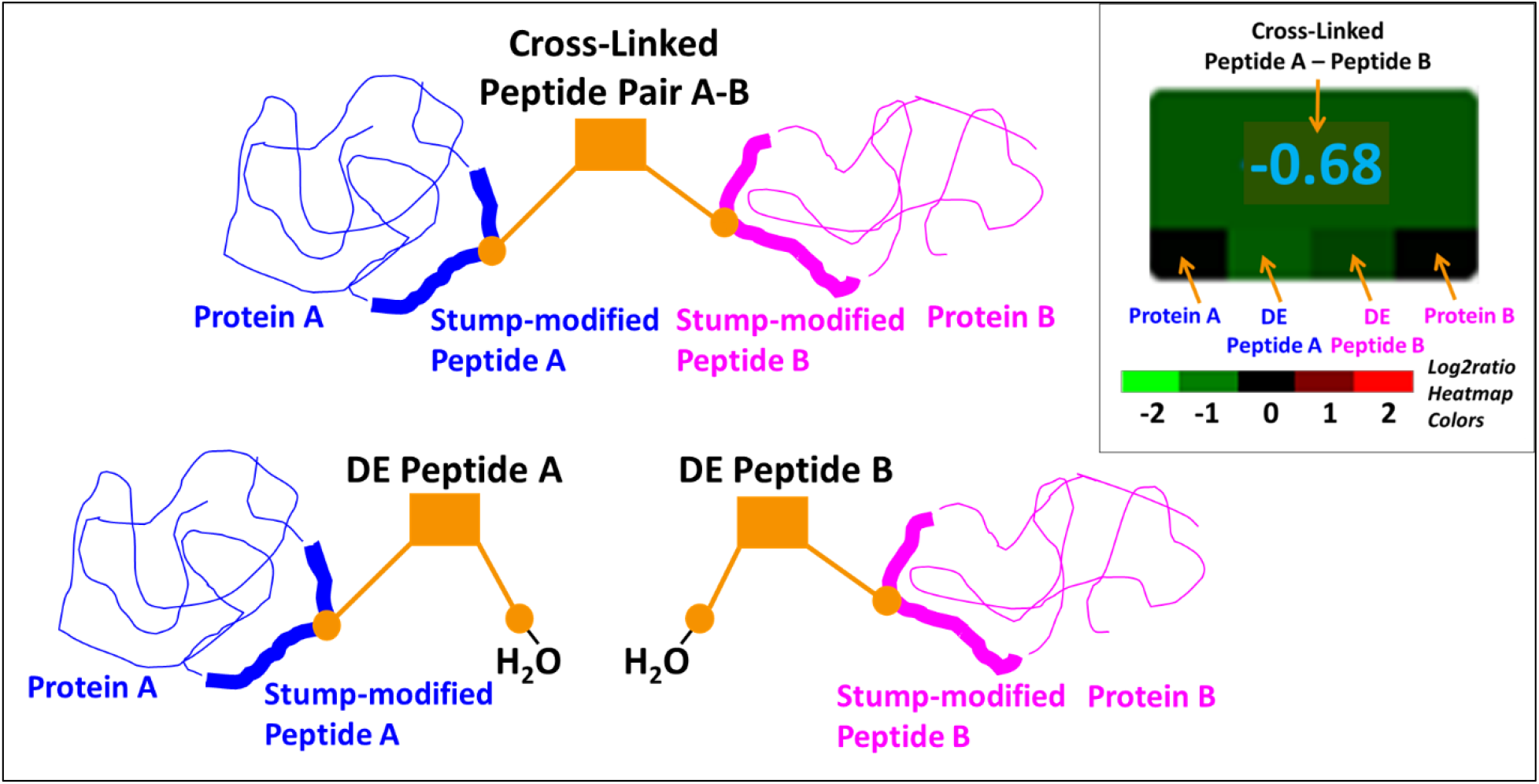
Display of cross-links alongside their corresponding DE peptides in XLinkDB. Cross-linked peptide pair A-B spanning proteins A and B cross-linked at residues with residual stump modifications shown as gold-colored circles and corresponding DE peptides A and B linked to the same protein residues of protein A and protein B, respectively. Shown in the inset (upper right) is the display in the XLinkDB color-scale heatmap of the quantitation of cross-links alongside quantitation of their corresponding proteins and DE peptides to facilitate comparison. The colors indicate each log2ratio value. In addition, the cross-link log2ratio (−0.68) is displayed in the heatmap while those of the proteins and DE peptides are shown in mouseover of their heatmap area.

### Facilitated interpretation of cross-link abundance level changes using corresponding DE peptides

For a cross-link with a log2ratio significantly different from zero, reflecting different abundance in the two compared samples that is not driven by protein abundance level changes, it is possible to observe two different common relationships between the cross-link and DE peptide log2ratios (Supporting Figure S1). In the first case, both DE peptides do not display altered abundance in the two samples despite the cross-link doing so. One can thus conclude that the observed cross-link abundance change is not due to conformational changes coinciding with altered solvent accessibility or reactivity of the cross-linked residues since this would also affect corresponding DE peptide levels. Nor is it due to altered PTMs on the cross-linked peptides or preceding lysines or arginines that might alter concentration of the tryptic peptides. Instead, the most likely explanation for the cross-link abundance change in such a case is an alteration in protein configuration that changes the solvent-accessible distance separating the cross-linked residues. In the case of an inter-protein cross-link, this could reflect altered protein interaction. Leveraging the DE quantitation is thus very useful since many possible explanations can be eliminated.

In the second case, one or both of the DE peptides have the same abundance change as the cross-link. The simplest interpretation of this situation is that the observed difference of cross-link abundance comes from a localized change in solvent accessibility or reactivity of the attached residue of the DE peptide with the same abundance change, or alteration of PTMs on that DE peptide or its preceding lysine or arginine. These scenarios would result in a similar change in abundance of the cross-link since one or more of its peptides would be altered in abundance like the DE peptide. A conformational change altering the solvent-accessible distance between the cross-linked residues is not required but could also occur, perhaps as a consequence or cause of the alterations localized at the DE residues. Leveraging the DE quantitation is again very useful since in this case it indicates a conformational change localized to a cross-linked residue.

XLinkDB uses Anova p-values to determine whether quantitation of a cross-link is similar to that of both cross-linked proteins from which it originated and with that of both corresponding DE peptides. Anova p-values are also used to assess whether DE peptide quantitation is similar to that of the protein from which it originated. A ‘hypothesis’ column is displayed in XLinkDB heatmap tables to indicate the simplest possible explanations for observed cross-link quantitation different from that of its cross-linked proteins on the basis of the corresponding DE peptide quantitation (see Methods). When none of the corresponding DE peptides have log2ratios significantly different from those of the originating protein, it is proposed that the abundance difference of the cross-link in the two compared samples is due to a conformational change altering the solvent-accessible distance between cross-linked residues. In contrast, when a DE peptide corresponding to a cross-link has a log2ratio that is similar to that of the cross-link, it is proposed that the observed cross-link abundance difference is possibly due to a conformational change localized to the DE peptide altering the solvent accessibility or reactivity of its attached residue or levels of PTMs on the peptide itself or reported by dbPTM^31,32^ to occur on residues of the peptide or its preceding amino acid. A mouseover of a displayed hypothesis indicates the calculated Anova p-values upon which it was based. One can filter the table for cross-links with agreeing or disagreeing corresponding DE peptide quantitation. With this functionality, researchers can easily identify cross-links with different abundance levels in two samples either likely due to conformational changes altering the solvent-accessible distance separating the cross-linked residues, or to more localized conformational differences at the cross-linked residues. The low variance of iqPIR quantified log2ratios helps enable this assessment, though XLinkDB supports all quantitative methods that calculate cross-link ratios in two compared samples^26^. A tutorial video explaining how to view quantitation of cross-links alongside their corresponding DE peptides in XLinkDB can be seen in **Supporting Information Video S1**.

### Integrative analysis of DE and protein interactome changes in TAC versus Sham mice

Previously described iqPIR cross-link data comparing mitochondrial protein interactomes in six replicate pairs of failing heart (TAC) versus healthy (Sham) mice^28^ was uploaded to XLinkDB along with DE peptides (dataset <u>Mito_iqPIR_TACsham_withDeadends_Bruce</u>). This dataset contains a total of 5,633 cross-links, 1,549 of which are looplinks, as well as 3,682 DE peptides. A total of 904 cross-links were quantified in at least 4/6 replicates with confident quantitation (See Methods). Of those, 374 (41%) have a single corresponding quantified DE peptide, and 279 (31%) have two. Cross-links with quantitation that differs significantly from that of its proteins of origin, as determined by label-free quantitation (LFQ), can be identified filtering by Anova p-value. In total, 286 such cross-links are present, of which 124 and 67 have one and two corresponding quantified DE peptides, respectively. Of the 124 cross-links with one corresponding DE peptide, 81 have similarly altered DE peptide abundance in TAC versus Sham, and 43 have DE peptides with no significant abundance difference. Of the 67 cross-links with two corresponding DE peptides, 46 have similarly differing DE peptide abundance in TAC versus Sham, 17 have DE peptides with no significant abundance difference, and 4 have one DE peptide of each type.

For inter-protein cross-links, including homodimers, consideration of the quantitation of corresponding DE peptides can indicate whether a change of cross-link abundance that is not driven by protein level changes is caused by changes in the solvent-accessible distance between the cross-linked residues, including association/dissociation, or whether perhaps it reflects other more localized conformational differences affecting the solvent accessibility or reactivity of the cross-linked residues. An inter-protein homodimer cross-link with increased abundance in TAC versus Sham (log2ratio 0.49) spans two protein molecules of AATM, mitochondrial aspartate aminotransferase, at residue 82 (Figure 2A). In contrast, the abundance of the DE peptide linked to the protein at residue 82, and that of the protein itself, do not differ between TAC and Sham mice. The hypothesis proposed to explain the observed increase in cross-link abundance is increased dimerization. This integrated analysis does not suggest that changes in solvent accessibility or reactivity of residue 82, or reported PTMs at that residue or residue 90, could explain the observed cross-link quantitation since that would have resulted in DE peptide abundance change. AATM dimerization is induced by substrates^33^, and its activity is elevated in patients with cardiovascular disease^34^. It is thus possible that the observed increase in dimerization is associated with increased activity of the protein in TAC versus Sham mice.

**Figure 2.**
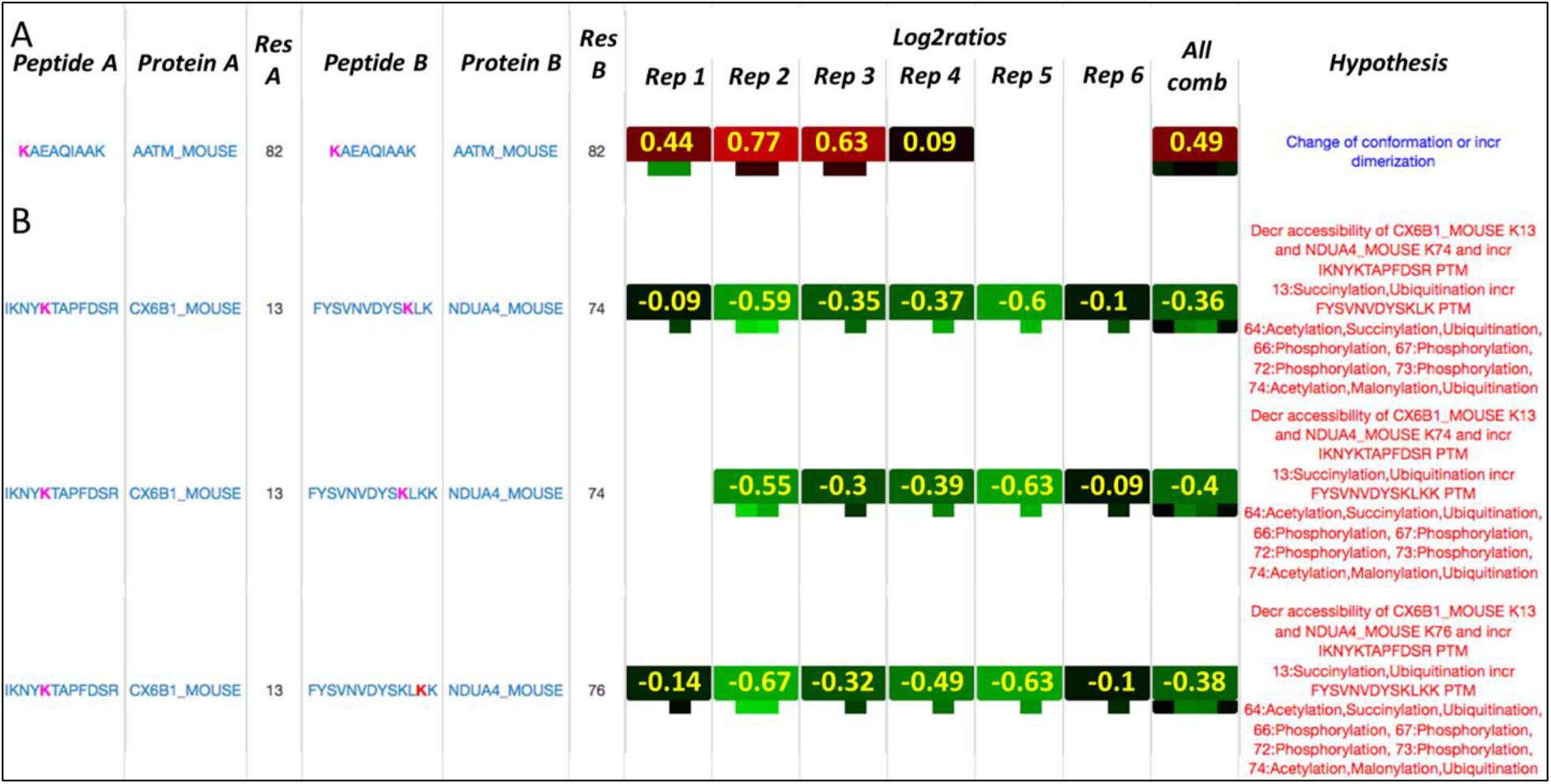
Four inter-protein cross-links with altered abundance in TAC versus Sham mice. A. Quantitation of homodimeric cross-link of AATM at residue 82 and its corresponding DE peptide in six paired TAC/Sham replicate mice and a combined ratio. Heatmap colors indicate log2ratio values ranging from green (−2) to black (0) to red (2). DE peptide at residue 82 displays unchanged abundance in TAC versus Sham. The displayed hypothesis includes increased dimerization. B. Three cross-links spanning CX6B1 and NDUA4 components of cytochrome c oxidase with decreased abundance in TAC versus Sham mice. Corresponding DE peptides at residues 13 of CX6B1 and 74 and 76 of NDUA4 exhibit similarly decreased abundance in TAC. The proposed hypothesis for the decreased abundance of all three cross-links is decreased solvent accessibility or reactivity of both cross-linked residues, or increased reported PTMs on both cross-linked peptides.

Three inter-protein cross-links spanning CX6B1 and NDUA4, two proteins of the mitochondrial cytochrome c oxidase component of Complex IV, which transfers two electrons to reduce oxygen into water, display decreased abundance in TAC versus Sham mice (log2ratios -0.4 to -0.36), and their corresponding DE peptides at CX6B1 residue 13 and NDUA4 residues 74 and 76 each display a similar change in abundance with log2ratios in the range of -0.57 to -0.36 (Figure 2B). In contrast, the levels of both proteins are not changed. The automatically generated hypotheses for the decreased cross-link abundances propose reduced accessibility of CX6B1 residue 13 and NDUA4 residues 74 and 76 or increased reported PTMs on CX6B1 peptide IKNY**K**^13^TAPFDSR residue 13 and NDUA4 peptide FYSVNVDYS**K**^74^LK residues 64, 66, 67, 72, 73, or 74. Complex IV activity was reduced in TAC mice^28^, and the observed change of interaction between CX6B1 and NDUA4 was proposed to possibly play a role. Therefore, it is of great interest that the quantitation of DE peptides indicates conformational changes of both proteins localized to their cross-linked residues. Future studies can target the indicated PTMs and assess abundance in TAC and Sham animals, particularly acetylation sites since that modification is known to be increased in TAC^35,36^. It is possible that the two proteins also dissociate in TAC, though that is not required to explain the cross-link abundance change. Assessment of this interaction level with conventional methods involving nondenaturing gel electrophoresis has been difficult due to the challenges associated with sensitivity of the NDUA4-Complex IV interaction to conditions for solubilization^37^.

For intra-protein cross-links, consideration of the quantitation of corresponding DE peptides can reveal whether a difference of cross-link abundance not driven by protein level changes reflects protein conformational changes altering the solvent-accessible distance separating the cross-linked residues or whether perhaps it reflects other more localized differences affecting the solvent accessibility or reactivity of those residues. Mitochondrial ADP/ATP carrier ADT1 is a dual-gated transporter shuttling ADP and ATP between the cytosol and mitochondrial matrix, alternating between the c-state conformation open to the intermembrane space and the m-state conformation open to the mitochondrial matrix. State-specific salt bridges are critical to stabilize the alternative channel conformations. Although its protein levels are not altered in TAC versus Sham mice, several of its intra-protein cross-links are increased in abundance. Because both c-state and m-state specific cross-links were increased in heart failure samples, an open channel AlphaFold model ADT1 conformation compatible with all increased cross-links was proposed to be increased in TAC mice^28^. This conformation is consistent with the previously proposed existence of a “P state” of ADT1 described by Karch *et al*.^38^ and Bround *et al*.^39^.

Figure 3A shows the log2ratios of four intra-protein cross-links of ADT1 with increased abundance in TAC versus Sham mice (log2ratios 0.67 – 0.86) along with quantitation of their corresponding DE peptides and protein. According to their Jwalk^40^ SASDs, the first cross-link is not consistent with the c-state conformation, while the last two cross-links are not consistent with the m-state conformation. It is immediately apparent based on quantitation of their corresponding DE peptides that the abundance change of the first three cross-links must involve localized conformational differences such as a change of solvent accessibility or reactivity of a cross-linked residue or altered peptide PTMs. The DE peptide linked at residue 33 corresponding to the first two cross-links increases in abundance (log2ratio 1.22). Since that residue participates in salt bridge formation that is critical for gate closure in the c-state conformation^41^, an increase in the abundance of the K33 DE peptide is consistent with increased reactivity due to a reduction in salt bridge formation and thus, a reduction of c-state gate closure. This change at residue 33 is sufficient to explain the observed cross-link abundance changes. Figure 3B shows the salt bridge at residue 33 present in the c-state conformation and absent in the m-state conformation. Interestingly, the second cross-link spanning residues 23-33 has a Jwalk SASD consistent with both the c-state and m-state conformations. Taking into consideration the expected reduced reactivity of residue 33 in the c-state due to salt bridge formation, however, one must conclude that the K23-K33 cross-link is less likely derived from the c-state conformation. The hypotheses automatically generated by XLinkDB for the first two cross-links on the basis of Anova p-values comparing cross-link, DE peptide, and protein quantitation suggests increased solvent accessibility at K33, consistent with reduced salt bridge formation. It also suggests the possibility of reduced reported PTMs at residues 33, 42, and 43 that would result in increased abundance of the tryptic peptide V**K**^33^LLLQVQHASK.

**Figure 3.**
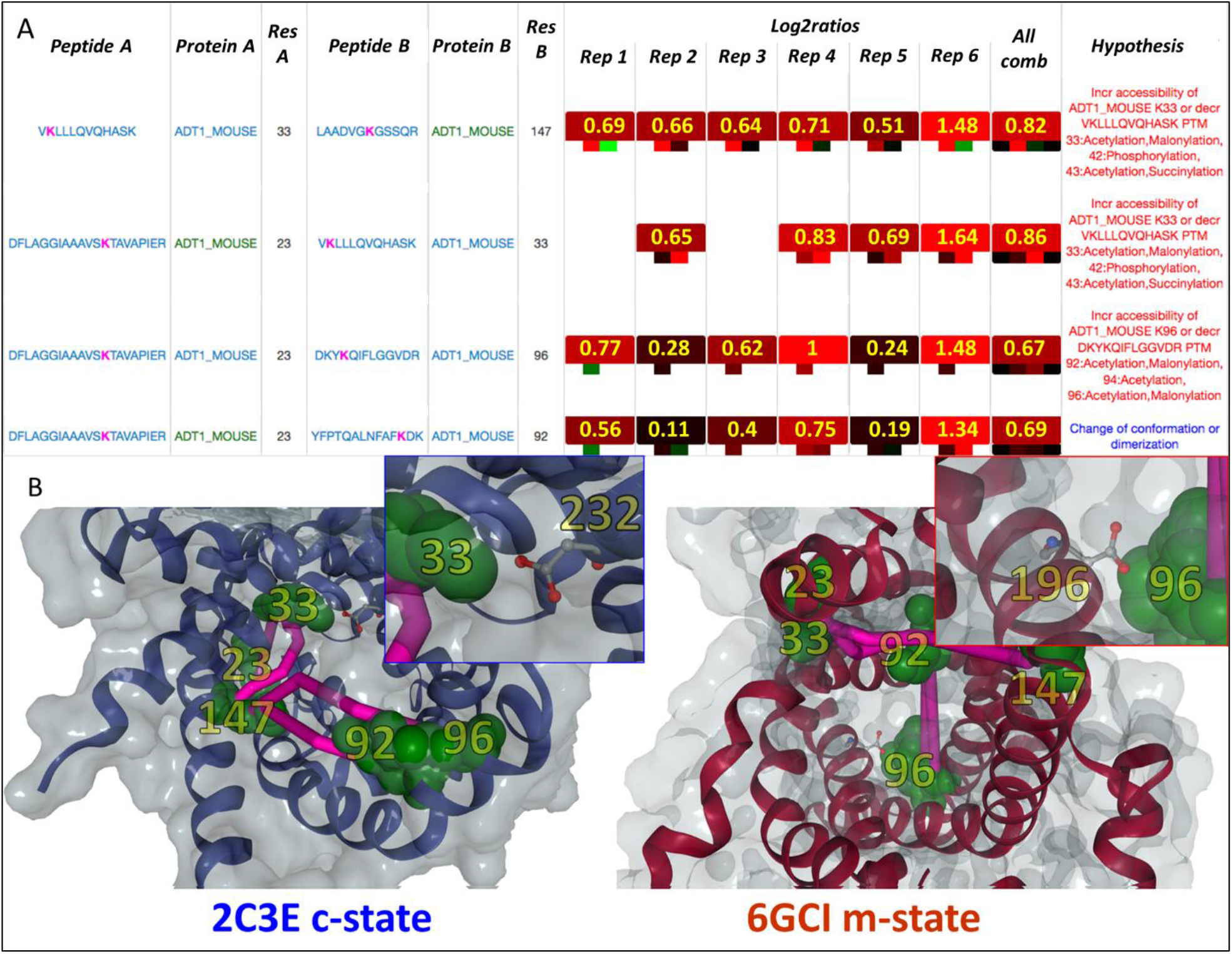
Four intra-protein cross-links of ADT1 with increased abundance in TAC versus Sham mice. A. Quantitation of cross-links and their corresponding DE peptides in six paired TAC/Sham replicate mice and a combined ratio. Heatmap colors indicate log2ratio values ranging from green (−2) to black (0) to red (2). DE peptides at residues 33 and 96 display similar increased TAC abundance. Hypotheses shown include increased reactivity of residues 33 and 96 consistent with removal of key salt bridges maintaining the c-state and m-state conformation, respectively. B. Cross-links shown in context of c-state and m-state structures with insets showing the salt bridges between residues K33 and D232 (c-state) and between residues K96 and D196 (m-state). Acidic residues are shown with ball and stick structures. Cross-linked lysine residues are depicted in green and the Jwalk solvent accessible path between them, in pink. Protein residue positions are shown in yellow text. For clarity, only one of the two copies of ADT1 (chain A) in 6GCI is shown.

In the case of the third cross-link (Figure 3A), the DE peptide linked at residue 96 increases in abundance (log2ratio 0.53) and is known to participate in salt bridge formation important for maintaining the m-state configuration^42^. The increase in DE peptide abundance is consistent with increased reactivity due to a reduction in the salt bridge formation and thus, a reduction of the m-state conformation in TAC mice. Figure 3B shows the salt bridge at residue 96 absent in the c-state conformation and present in the m-state conformation. The hypothesis generated by XLinkDB suggests increased solvent accessibility at residue 96, consistent with reduced salt bridge formation. It also suggests the possibility of reduced reported PTMs at residues 92, 94, and 96 that would result in increased abundance of the tryptic peptide DKY**K**^96^QIFLGGVDR.

Both DE peptides corresponding to the fourth cross-link spanning residues 23-92 have unchanged abundance in TAC versus Sham suggesting that the increase in cross-link abundance is solely due to a change of conformation altering the solvent-accessible distance separating the cross-linked residues, without any localized changes in solvent accessibility or reactivity of those residues. The hypothesis generated by XLinkDB on the basis of Anova p-values comparing cross-link, DE peptide, and protein quantitation suggests a conformational change. In this case, it is not likely that reduced known PTMs at residues 10, 22, 23, or 24 of tryptic peptide DFLAGGIAAAVS**K**^23^TAVAPIER or at residues 81, 84, 92, or 94 of tryptic peptide YFPTQALNFAF**K**^92^DK could be responsible for the observed increase in cross-link abundance. Since one structure of ADT1, 6GCI, contains a protein dimer, it is also suggested that the cross-link abundance change could be due to altered dimerization of the protein.

Because cross-links thought to be specific to both the m-state and c-state were found to be increased in abundance in TAC versus Sham mice, it was proposed that there is a third open channel conformation consistent with all the cross-links that is increased in TAC mice^28^. The quantitation of DE peptides at the state-specific salt bridge residues 33 and 96 further suggests that there is a reduction in both c-state and m-state conformations. The hypotheses generated automatically by XLinkDB to explain the observed increased abundances of the four cross-links in TAC versus Sham mice include a change of conformation of the protein coinciding with increased reactivity at residues 33 and 96. This is consistent with diminished c-state and m-state conformations of ADT1 in TAC mice as a consequence of an increase in the open channel configuration. Such a change would yield increased ADT1-mediated proton leak^43^ through the mitochondrial inner membrane and consequently, decreased membrane potential available to motivate ATP production in heart failure.

## Conclusions

XLinkDB now supports DE peptides, enabling the display of quantitation of cross-links alongside the quantitation of their corresponding DE peptides and proteins. For the first time, one can leverage quantitation of DE peptides to easily assess whether the abundance change of a cross-link is likely due to a localized conformational change affecting the solvent accessibility or reactivity of the cross-linked residue or levels of peptide PTMs. We have demonstrated how this combined information helps to reveal underlying conformational causes of observed cross-link abundance differences between two samples that are not attributable to protein level changes. For example, the intra-protein cross-link of ADT1 spanning residues 23-33 exhibited increased levels in TAC versus Sham mice. This cross-link had a Jwalk SASD consistent with both the c-state and m-state conformations of ADT1, so it wasn’t clear whether its increased abundance might be due to increased levels of either or both states. However, because the corresponding DE peptide attached at residue 33 had similarly increased abundance, one could consider the increased levels of the cross-link as possibly being due to increased solvent accessibility or reactivity of that residue. K33 is known to be involved in salt-bridge formation critical to maintaining the c-state conformation of ADT1, so its increased accessibility or reactivity in TAC is consistent with reduced salt-bridge formation, and hence, reduced levels of the c-state conformation. This explanation for the increased abundance of the cross-link was not apparent from the cross-link quantitation on its own.

XLinkDB uses Anova p-values to compare the cross-link, DE peptide, and protein quantitation in order to display in a ‘hypothesis’ column possible explanations for observed cross-link abundance differences not due to protein level change; explanations that are consistent with the combined observed quantitation based on current understanding of conformational features that influence cross-link and DE peptide abundance. In cases in which the corresponding DE peptides do not have differing abundances, one can attribute the cross-link abundance difference to an altered conformation affecting the solvent-accessible distance between the cross-linked residues, including in the case of inter-protein cross-links, increased association or dissociation of the two spanned proteins. This is the interpretation usually given by default in the absence of corresponding DE peptide quantitation. In contrast, when one or both corresponding DE peptides exhibit abundance differences similar to that of the cross-link, this observation is most likely caused by a localized conformational change affecting the solvent accessibility or reactivity of that DE peptide residue or by a change in PTMs residing on residues of the peptide. It is still possible that an altered conformation affecting the solvent-accessible distance between the cross-linked residues also occurs coincident with those changes. Additional more complicated explanations for observed cross-link abundance differences are undoubtedly also possible so the explanations displayed in the XLinkDB hypothesis column must be used only as a guide. We showed how the proposed explanations for the ATD1 intra-protein cross-links with increased abundance in TAC versus Sham mice matched the previously published conclusions of scientists that required much time and effort to survey the data and literature. Researchers can assess the proposed explanations consistent with their cross-link and DE peptide quantitation and then direct future investigations to pursue any of interest. For example, if a change in abundance of certain PTMs is proposed as possibly responsible for the cross-link abundance difference, one can direct experiments to detect and quantify those modifications in the compared samples.

If a cross-linked peptide contains PTMs, then a change in the abundance of those PTMs in the two compared samples would be directly reflected in the abundance of that peptide. In addition, a change of abundance of any other PTMs, not present on the peptide but nevertheless occurring on its residues or adjacent enzyme cleavage sites, can inversely influence the abundance of the peptide. Although a high occupancy of such modifications in at least one of the two compared samples may be required for detection as a cross-linked peptide abundance difference, all PTMs occurring on the peptide residues, not just a single one, contribute. For example, there are a wide variety of PTMs on lysine residues so it is possible that an increase in several, such as acetylation and succinylation, and at multiple lysines in a cross-linked peptide, could collectively result in an observed reduced peptide abundance.

Analyzing DE peptides together with cross-links adds value to quantitative cross-link data. We are currently seeking ways to optimize the acquisition of DE peptides to generate as much coverage of cross-links as possible. Ideally, it would be beneficial if the majority of cross-linked peptides that are identified and quantified are also identified and quantified as DE peptides. We have focused initially on cross-links that have different abundances in two samples under comparison despite the cross-linked protein levels not differing, since they are the most challenging to interpret. In the future, we can include in the displayed hypothesis column cases in which the difference in cross-link abundance could be attributable to protein level changes. We will continually assess whether other relationships between cross-link and DE peptide quantitation are observed, such as abundance changes in opposite directions, and seek possible explanations for them. Researchers can leverage this functionality by uploading their quantitative cross-link data, including DE peptides, to XLinkDB.

## SUPPORTING INFORMATION

The following supporting information is available free of charge at ACS website

- Figure S1 – Comparison of combined replicate log2ratios in TAC versus Sham mice calculated for cross-links and corresponding DE peptides.
- Video S1 – Training tutorial on how to view on XLinkDB quantitation of cross-links alongside quantitation of their corresponding DE peptides and displayed possible explanations for observed cross-link abundance differences.

## Acknowledgements

The authors acknowledge and thank all members of the Bruce lab for helpful comments and suggestions during the course of preparation of this manuscript. This work was supported by the following grants from the National Institutes of Health: R35GM136255, R01AG078279, R56AG078279, R56AG070096 and R01HL144778.

