## Supplemental Figure S1 for "Integrated Analysis of Cross-Links and Dead-End Peptides for Enhanced Interpretation of Quantitative XL-MS"

**Supporting Information**

**Table of Contents** (list of supplementary components)

- Figure S1 – Comparison of combined replicate log2ratios in TAC versus Sham mice calculated for cross-links and corresponding DE peptides.

- Video S1 – Training tutorial on how to view on XLinkDB quantitation of cross-links alongside quantitation of their corresponding DE peptides and displayed possible explanations for observed cross-link abundance differences.

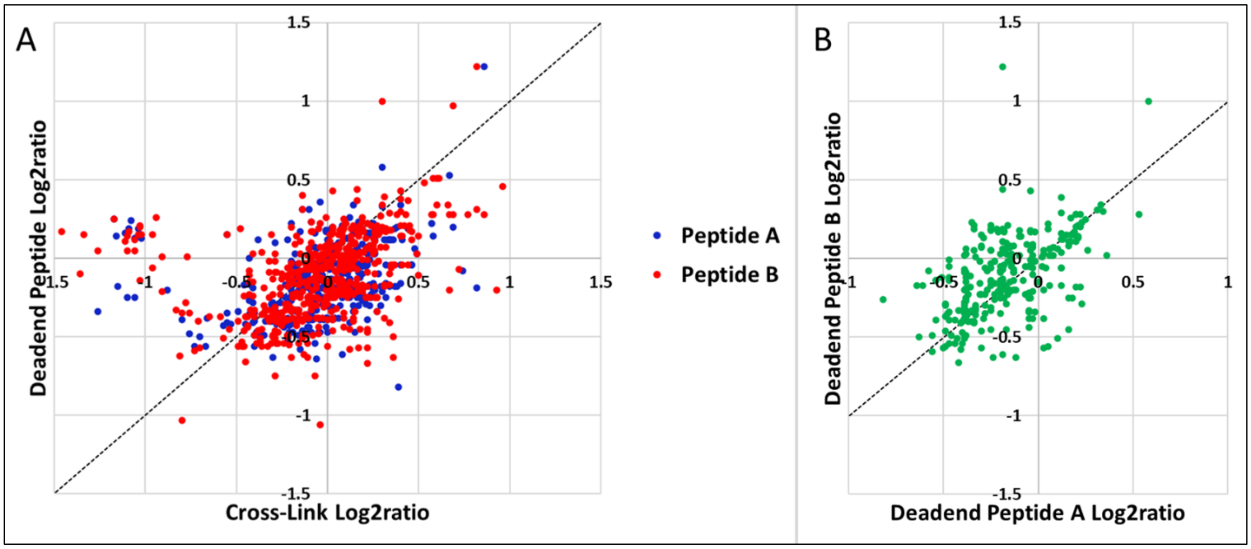


Supporting Figure S1. **Comparison of combined replicate log2ratios in TAC versus Sham mice calculated for cross-links and corresponding DE peptides.** A. Cross-linked Peptide A-B versus corresponding DE Peptides A and B. Cases along the 45 degree line far away from the origin are ones in which the cross-link abundance difference in TAC versus Sham mice is due, at least in part, to a conformational change localized to the DE peptides. B. DE peptide A versus DE peptide B. Each pair corresponds to cross-linked peptide A-B.
